# Design, Synthesis, and Efficacy of Novel Trojan Horse Compounds Targeting Metabolic Vulnerabilities in Prostate Cancer

**DOI:** 10.1101/2024.08.29.610353

**Authors:** L Bogue Edgerton, E F Hayball, F O’Connell, MP Ward, S O’Toole, CM Martin, S Selemidis, RD Brooks, P Tewari, DA Brooks, JJ O’Leary

## Abstract

**Background and purpose:** Prostate cancer (PCa) remains a significant global health concern, warranting the development of new therapeutic strategies.^1^ Metabolic reprogramming, an emerging hallmark of PCa progression characterized by aberrant nutrient utilization, has become a focus area in developing new treatments. Among these metabolic alterations, the Warburg effect—a shift towards aerobic glycolysis— has garnered significant attention as a potential therapeutic target. ^2^ To explore this, we synthesised a novel class of Trojan Horse compounds designed to exploit the unique metabolic profile of cancer cells and investigated their therapeutic potential using cell based models of PCa, as a proof of principle strategy.

**Experimental approach:** Preliminary cytotoxicity analysis revealed that native menadione had low efficacy against PCa cells. Consequently, we developed glucose and fatty acid conjugated TH compounds based on a menadione backbone. A library of six novel menadione-glucose and fatty acid-based TH compounds were synthesized through copper-catalysed click chemistry. These compounds were designed to mimic a desirable fuel source (sugar or fatty acid) acting as a “Trojan Horse” for PCa cells to modulate their metabolism. The cytotoxic and selective effects of the TH compounds were evaluated in PCa and non-malignant cell lines cultured under varying glucose conditions. To elucidate the mechanisms of action, mitochondrial bioenergetics, metabolic phenotyping, and metabolomic profiling were employed to assess the impact of each TH compound on cellular metabolism.

**Key results:** TH3 (glucose) and TH5 (fatty acid) compounds demonstrated promising cytotoxicity for PCa cell lines, with TH5 exhibiting superior selectivity compared to TH3 and particularly over native menadione. Despite these results, alterations in metabolic phenotypes were only modest. Mitochondrial bioenergetics were notably impacted by TH5, while ROS levels remained stable post-treatment, likely due to an increase in ROS scavenging amino acids as a compensatory antioxidant response.

**Conclusion and Implications:** This study demonstrates the potential of employing glucose and fatty acid-conjugated compounds to target the unique metabolic features of PCa. These findings offer promising new avenues for further investigation, aimed at optimizing efficacy and selectivity of these compounds for clinical translation. The intricate relationship between ROS production and antioxidant defence mechanisms emphasizes the complex nature of metabolic interventions in cancer. Elucidating the mode of action and exploring molecular modifications may pave the way for personalised and targeted PCa therapies, ultimately improving patient outcomes and mitigating the global burden of this disease.

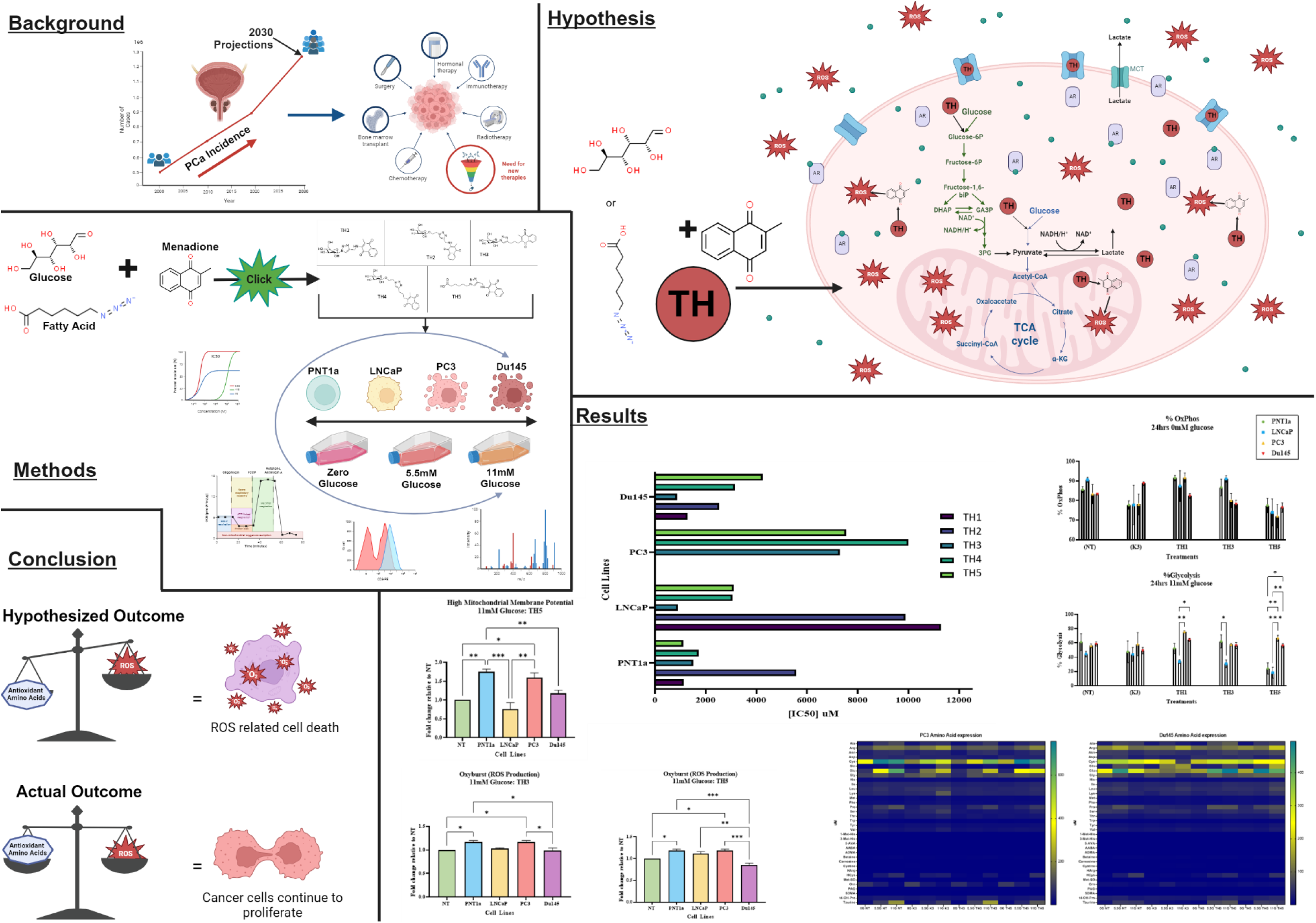

**Significance statement:** This body of work gives critical insights into the metabolic vulnerabilities of prostate cancer (PCa), particularly of androgen-independent and metastatic disease, which currently present significant treatment challenges. By designing, synthesising and testing innovative menadione-based ‘Trojan Horse’ compounds that target altered cancer cell metabolism, this research highlights a promising new therapeutic strategy against malignant cells. The findings underscore the potential for metabolic reprogramming to combat treatment resistance in advanced PCa, paving the way forward for the development of new drugs and personalised treatment approaches. This could lead to improved outcomes for patients with currently untreatable forms of the disease.

## 1.0 Introduction

Prostate cancer (PCa) is the second most prevalent cancer among men globally, with its incidence predicted to double by 2030. ^3, 4^ The growth and survival of early-stage PCa are dependent on androgen hormones, and therapeutic approaches targeting androgen cycles, such as androgen ablation therapy (AAT) are in clinical practice. ^5,6^ However, advanced-stage, androgen-independent prostate cancers (AIPCs) pose a significant challenge for treatment, as they are often incurable with current therapies.^7–10^ To address this significant health challenge, we proposed a novel approach that targets the unique metabolic pathways of PCa, which are androgen-independent and thus are not amenable to AAT. Given the ever-increasing incidence and mortality of cancer, this novel approach is important. Our research has developed a unique approach which focuses on targeting the cancer cell’s altered metabolic pathway, using ‘Trojan horse’ compounds. The overarching approach is to deliver toxic reactive oxygen species (ROS) via menadione, a synthetic analogue of vitamin K3, which is complexed to simple sugars and lipids to specifically target and kill cancer cells while sparing non-malignant cells. This approach builds on existing studies into Vitamin K_3_ ‘s effects on PCa^11^ and leveraging menadione’s ROS-dependent cytotoxicity seen in various cancer cell types.^12,13,14^ Previous phase I clinical trials have shown safety of menadione, both alone and in combination with other treatments,^15^ and phase II trials have highlighted potential efficacy.^16^

When targeting PCa metabolism with metabolic substrates, it is important to note that the normal prostate has a distinct metabolic signature characterised by zinc accumulation. This zinc presence inhibits mitochondrial aconitase to limit Krebs cycle activity.^17,18^ In contrast, cancer cells, including PCa, undergo metabolic reprogramming, a hallmark of malignancy leading to an increased reliance on cytosolic metabolism, such as glycolysis.^19,20^ PCa cells, in particular, display a shift towards glycolysis, even in the presence of oxygen, despite a decreased efficiency in glucose metabolism.^17,21,22^ This metabolic shift prioritizes faster energy production, to support the demands of the heightened and sustained proliferation of cancer cells, albeit at the cost of generating excessive metabolic waste.^23^

Glucose plays a critical role in cancer cell metabolism, serving as a primary substrate for ATP production and biomass synthesis. Its availability is a key factor influencing cancer cell metabolism, therapeutic response, and cellular redox balance.^24,25^ High glucose availability can promote an increase in glycolytic flux, resulting in sustained proliferation and cell survival. However, increased glucose availability may reduce cancer cell sensitivity to therapeutic intervention.^26^ In terms of redox status, high glucose levels are thought to promote the production of ROS, resulting in oxidative damage and potentially promoting cancer progression.^27,28^ Alternatively, glucose deprivation is thought to induce metabolic stress, which could sensitise cancer cells to treatment by altering their ability to generate ATP and repair DNA damage while also increasing cancer cell vulnerability to oxidative stress-induced damage or death.^29–31^

Understanding the dynamic interplay between energy metabolism, mitochondrial function, and redox signalling in PCa is important for developing effective therapeutic strategies that target metabolism and limit cancer cell adaption.^32^ Therefore the aim of the present study was to synthesise novel menadione-based TH compounds using click chemistry to specifically target the altered metabolism of cancer cells. Our findings highlight the potential of metabolic targeting strategies to disrupt PCa metabolism and enhance the efficacy of both new and existing therapies.^2^

## 2.0 Materials and Methods

### 2.1 Materials

#### Cell Lines and Media

PNT1a (Sigma-Aldrich), LNCaP (ATCC, Manassas, VA, USA), PC3 (ATCC), Du145 (ATCC), RPMI-1640 media (Sigma-Aldrich), Glucose Free RPMI-1640 media (Sigma-Aldrich), Dialyzed foetal bovine serum (Sigma-Aldrich), Penicillin-streptomycin (Sigma-Aldrich).

#### Synthesis Reagents

Merck (Sigma Aldrich) - Menadione, propargylamine, 5-hexynoic acid, 6-azidohexanoic acid, copper(II) sulfate pentahydrate (CuSO_4_·5H_2_O), (+)-sodium-L-ascorbate, 5-hexynoic acid, ammonium persulfate ((NH_4_)_2_S_2_O_8_), silver nitrate (AgNO_3_), methanol (MeOH), dichloromethane (CH_2_Cl_2_), dimethyl sulfoxide, 20–63 µm grade silica gel, silica gel 60 F_254_ aluminium plates, ethyl acetate (EtOAc), hexane, dimethyl formamide (DMF). Synthose. Inc - β-D-Glucopyranosyl azide, 2-Azidoethyl β-D-Glucopyranoside. ChemSupply Australia - ethanol (EtOH), sodium hydrogen carbonate, sodium chloride, sodium sulfate (Na_2_SO_4_) anhydrous.

#### Cytotoxicity and Selectivity Assay Reagents

Alamar blue assay kit (ThermoFisher), Dimethyl sulfoxide (DMSO) (Sigma-Aldrich)

#### Metabolic Phenotype and Bioenergetics Assay Reagents

Oligomycin A (Sigma-Aldrich), FCCP (Sigma-Aldrich), Antimycin A (Sigma-Aldrich), XF assay media (Agilent Technologies, Santa Clara, CA, USA), Crystal violet (Sigma-Aldrich)

#### ROS and MMP Flow Cytometry Reagents

Fluorescent dyes (Sigma-Aldrich and Thermofisher), FACS buffer (Thermofisher), Compensation beads (Thermofisher)

### 2.2 Cell Culture

This study utilized four prostate cell lines for drug testing including: PNT1a (Sigma-Aldrich), LNCaP (ATCC, Manassas, VA, USA), PC3 (ATCC), and Du145 (ATCC). The PNT1a cell line is immortalized with SV40, representing non-malignant human post-pubertal prostate cells. LNCaP, originated from a prostate supraclavicular lymph node metastasis is characterized by androgen dependency. PC3 was derived from prostate bone metastasis, while Du145 originated from a prostate brain metastasis, with both manifesting androgen independence.

All the cell lines were cultured under standard conditions in Roswell Park Memorial Institute (RPMI)-1640 media (Sigma–Aldrich), supplemented with 10% dialyzed foetal bovine serum (Sigma–Aldrich) and 5 mM penicillin-streptomycin (Sigma–Aldrich). They were maintained at 37°C in a humidified atmosphere containing 5% CO2.

### 2.3 Compound Synthesis

#### 2.3.1 General Methods

All chemicals purchased from commercial suppliers were used without further purification, except for anhydrous solvents, which were dried over molecular 3Å sieves. Column chromatography was performed using 20–63 µm grade silica gel. Thin layer chromatography was performed using aluminium sheets coated with silica gel 60 F_254_ purchased from Merck (Australia) and visualised using UV light (λ = 254 nm) and/or a KMnO_4_ oxidizing stain (KMnO_4_, K_2_CO_3_ and H_2_O). All ^1^H and ^13^C NMR spectra were collected on a BRUKER AVANCE III 500 MHz FT-NMR spectrometer. All NMR experiments were performed at 25 °C. Samples were dissolved in either CDCl_3_, DMSO-*d*_6_ or D_2_O, with the residual solvent peak used as the internal reference— CDCl_3_: 7.26 (^1^H) and 77.0 (^13^C); DMSO-*d*_6_: 2.50 (^1^H) and 39.52 (^13^C); D_2_O: 4.79 (^1^H). NMR multiplicities are reported as broad (br), singlet (s), doublet (d), triplet (t), quartet (q), pentet (p) and multiplet (m). All *J*-values are rounded to the nearest 0.1 Hz. High resolution mass spectrometry (HRMS) data was collected using an AB SCIEX TripleTOF 5600 mass spectrometer using a 95% MeOH in H_2_O mobile phase containing 0.1% formic acid.

#### 2.3.2 General CuAAC Procedure for Synthesis of TH Compounds

To the alkyne (1.2 equiv.) in DMF (0.2 molL^-1^) at room temperature (rt) protected from light was added a yellow solution of a copper(II) sulfate pentahydrate (0.5 equiv.) and (+)-sodium-L-ascorbate (1.0 equiv.) in 25% H_2_O in DMF (0.1 molL^-1^). The solution was stirred at rt for 1 min before addition of the azide (1.0 equiv.). The reaction was stirred at rt under air until the azide was consumed as indicated by TLC analysis (3 - 24 h). The reaction mixture was concentrated under a stream of N_2_ and the crude material was purified using flash chromatography (FC) on SiO_2_ (solvent system as specified) to give the triazole as specified (61 - 80%). The product was then dissolved in H_2_O, cooled to -80 °C overnight and lyophilized.

#### 2.3.3 2-(but-3-yn-1-ylamino)-3-methylnaphthalene-1,4-dione (3)

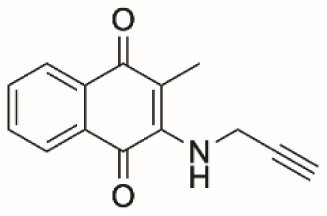

To a solution of menadione (1.47 g, 8.45 mmol, 1.0 equiv.) in EtOH (85 mL, 0.1 molL^-1^) protected from light was added propargylamine (1.09 mL, 17.1 mmol, 2.0 equiv.) at rt under a N_2_ atmosphere. The reaction was stirred at rt for 12 h and concentrated *in vacuo*. The crude dark red solid was purified using FC on SiO_2_ (20% EtOAc in *n*-hexane) to afford the product as orange needle-like crystals (1.13 g, 59%). Data for product matched that previously obtained in the literature.^33^

^1^H NMR (500 MHz, CDCl_3_) δ 8.08 (d, *J* = 7.6 Hz, 1H), 8.01 (d, *J* = 7.7 Hz, 1H), 7.68 (dd, *J* = 7.6, 7.6 Hz, 1H), 7.60 (dd, *J* = 7.5, 7.5 Hz, 1H), 5.78 (br s, 1H), 4.27 (dd, *J* = 6.7, 2.5 Hz, 2H), 2.35 (t, *J* = 2.5 Hz, 1H), 2.27 (s, 3H).

^13^C NMR (126 MHz, CDCl_3_) δ 184.0, 182.2, 145.5, 134.4, 133.2, 132.3, 130.4, 126.4, 126.2, 115.5, 80.1, 73.2, 35.3, 11.0.

HRMS (ESI-TOF, *m/z*) for C_14_H_12_NO_2_ 226.0790 [M + H]^+^, found 226.0220.

#### 2.3.4 2-methyl-3-(((1-((2*R*,3*R*,4*S*,5*S*,6*R*)-3,4,5-trihydroxy-6-(hydroxymethyl)tetrahydro-2*H*-pyran-2-yl)-1*H*-1,2,3-triazol-4-yl)methyl)amino)naphthalene-1,4-dione (Trojan Horse 1)

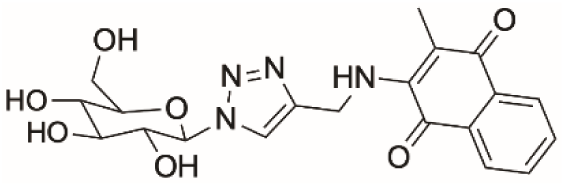

Prepared according to the General CuAAC procedure from amino alkyne **3** (146 mg, 0.648 mmol) and β-D-glucopyranosyl azide (111 mg, 0.541 mmol) (24 h, 5 to 10% MeOH in CH_2_Cl_2_) to afford the title compound as a red solid (182 mg, 78%).

^1^H NMR (500 MHz, D_2_O) δ 8.17 (s, 1H), 7.32 – 7.21 (m, 4H), 5.74 (d, *J* = 9.2 Hz, 1H), 4.57 (s, 2H), 4.02 (t, *J* = 9.2 Hz, 1H), 3.90 – 3.84 (m, 1H), 3.78 – 3.70 (m, 3H), 3.63 (t, *J* = 9.2 Hz, 1H), 1.59 (s, 3H).

^13^C NMR (126 MHz, D_2_O) δ 184.1, 182.0, 146.5, 146.3, 134.7, 132.5, 131.9, 129.5,

125.9, 125.3, 123.0, 112.6, 87.5, 78.8, 75.9, 72.3, 68.9, 60.4, 39.7, 9.8.

HRMS (ESI-TOF, *m/z*) for C_20_H_23_N_4_O_7_ 431.1488 [M + H]^+^, found 431.1434.

#### 2.3.5 2-methyl-3-(((1-(2-(((2*R*,3*R*,4*S*,5*S*,6*R*)-3,4,5-trihydroxy-6-(hydroxymethyl)tetrahydro-2*H*-pyran-2-yl)oxy)ethyl)-1*H*-1,2,3-triazol-4-yl)methyl)amino)naphthalene-1,4-dione (Trojan Horse 2)

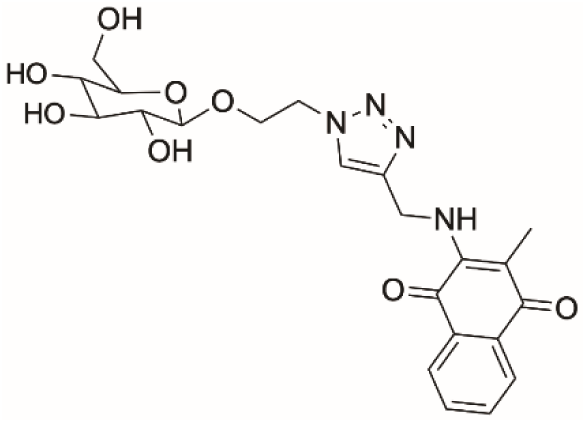

Prepared according to the General CuAAC procedure from the amino alkyne **3** (91 mg, 0.404 mmol) and 2-azidoethyl β-D-glucopyranoside (84 mg, 0.337 mmol) (18 h, 100% CH_2_Cl_2_ to 3% MeOH in CH_2_Cl_2_) to afford the title compound as a red solid (121 mg, 76%).

^1^H NMR (500 MHz, DMSO-*d*_6_) δ 8.06 (s, 1H), 7.93 (d, *J* = 7.6 Hz, 2H), 7.78 (app. t, *J* = 7.5 Hz, 1H), 7.69 (app. t, *J* = 7.6 Hz, 1H), 7.01 (t, *J* = 6.7 Hz, 1H), 4.81 (d, *J* = 6.4 Hz, 2H), 4.53 (t, *J* = 5.4 Hz, 2H), 4.19 (d, *J* = 7.7 Hz, 1H), 4.05 (dt, *J* = 10.9, 5.2 Hz, 1H), 3.87 (dt, *J* = 11.2, 5.5 Hz, 1H), 3.42 – 3.34 (m, 1H), 3.17 – 3.07 (m, 3H), 3.01 (t, *J* = 9.1 Hz, 1H), 2.93 (t, *J* = 8.3 Hz, 1H), 2.11 (s, 3H).

^13^C NMR (126 MHz, DMSO-*d*_6_) δ 181.9 (one obscured peak), 165.9, 146.5, 145.4, 134.4, 132.6, 132.3, 130.4, 125.7, 125.4, 123.5, 103.0, 77.1, 76.6, 73.3, 70.0, 67.4, 61.1, 49.7, 10.5.

HRMS (ESI-TOF, *m/z*) for C_22_H_27_N_4_O_8_ 475.1751 [M + H]^+^, found 475.1834.

#### 2.3.6 6-(4-(((3-methyl-1,4-dioxo-1,4-dihydronaphthalen-2-yl)amino)methyl)-1*H*-1,2,3-triazol-1-yl)hexanoic acid (Trojan Horse 5)

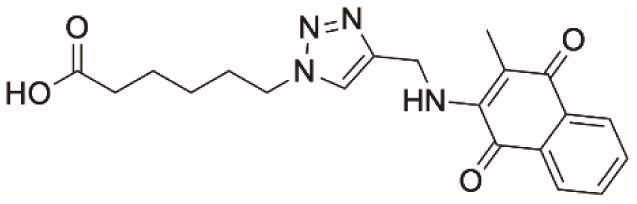

Prepared according to the General CuAAC procedure from amino alkyne **3** (76 mg, 0.338 mmol) and 6-azidohexanoic acid (45 µL, 0.307 mmol) (19 h, 50% EtOAc in *n*-hexane) to afford the product as a red solid (83 mg, 71%).

^1^H NMR (500 MHz, DMSO-*d*_6_) δ 7.98 (s, 1H), 7.92 (d, *J* = 7.6 Hz, 2H), 7.77 (dd, *J* = 7.6, 1.4 Hz, 1H), 7.69 (dd, *J* = 7.5, 1.3 Hz, 1H), 7.04 (t, *J* = 6.8 Hz, 1H), 4.80 (d, *J* = 6.7 Hz, 2H), 4.29 (t, *J* = 7.1 Hz, 2H), 2.14 (t, *J* = 7.4 Hz, 2H), 2.08 (s, 3H), 1.76 (p, *J* = 7.2 Hz, 2H), 1.47 (p, *J* = 7.5 Hz, 2H), 1.25 – 1.13 (m, 2H).

^13^C NMR (126 MHz, DMSO-*d*_6_) δ 182.1, 181.9, 174.3, 146.4, 145.7, 134.4, 132.6, 132.2, 130.4, 125.6, 125.4, 122.5, 112.1, 49.2, 33.4, 29.5, 25.4, 23.8, 10.4.

HRMS (ESI-TOF, *m/z*) for C_20_H_23_N_4_O_4_ 381.1641 [M - H]^-^, found 381.1982.

#### 2.3.7 2-methyl-3-(pent-4-yn-1-yl)naphthalene-1,4-dione (5)

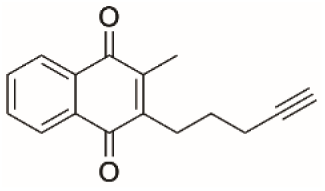

To menadione (500 mg, 2.90 mmol, 1 equiv.), 5-hexynoic acid (1.60 mL,14.5 mmol, 5.0 equiv.) and silver nitrate (493 mg, 2.90 mmol, 1.0 equiv.) in DMSO (30 mL) and H_2_O (300 µL) at rt protected from light was added ammonium persulfate (795 mg, 3.48 mmol, 1.2 equiv.). The reaction was stirred at 40 °C under a N_2_ atm for 17 h. The reaction was diluted with CH_2_Cl_2_ (10 mL) and the mixture was washed with sat. sodium bicarbonate solution (20 mL). The aqueous layer was extracted with CH_2_Cl_2_ (3 x 20 mL) and the combined organic extracts were washed with brine (70 mL), dried over anhydrous Na_2_SO_4_, filtered, and concentrated *in vacuo.* The crude product was purified using FC on SiO_2_ (100% CH_2_Cl_2_) to afford a yellow solid (360 mg, 52%). Data for product matched that previously obtained in the literature.^34^

^1^H NMR (500 MHz, CDCl_3_) δ 8.08 – 8.03 (m, 2H), 7.69 – 7.66 (m, 2H), 2.77 – 2.74 (m, 2H), 2.29 (td, *J* = 6.9, 2.6 Hz, 2H), 2.21 (s, 3H), 2.00 (t, *J* = 2.7 Hz, 1H), 1.76 – 1.65 (m, 2H).

^13^C NMR (126 MHz, CDCl_3_) δ 185.3, 184.7, 146.5, 143.9, 133.52, 133.50, 132.3 (2 overlapping peaks), 126.40, 126.36, 83.9, 69.2, 27.5, 26.3, 18.8, 12.8.

HRMS (ESI-TOF, *m/z*) for C_16_H_15_O_2_ 239.0994 [M + H]^+^, found 239.3654.

#### 2.3.8 2-methyl-3-(3-(1-((2*R*,3*R*,4*S*,5*S*,6*R*)-3,4,5-trihydroxy-6-(hydroxymethyl)tetrahydro-2*H*-pyran-2-yl)-1*H*-1,2,3-triazol-4-yl)propyl)naphthalene-1,4-dione (Trojan Horse 3)

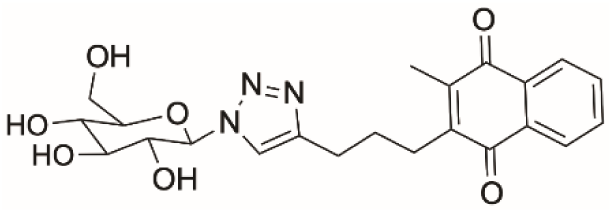

Prepared according to the General CuAAC procedure from alkyne **5** (115 mg, 0.508 mmol) and β-D-glucopyranosyl azide (95 mg, 0.462 mmol) (16 h, 5% MeOH in CH_2_Cl_2_) to afford the product as a yellow solid (163 mg, 80%).

^1^H NMR (500 MHz, DMSO-*d*_6_) δ 8.07 (s, 1H), 8.01 – 7.98 (m, 2H), 7.84 – 7.82 (m, 2H), 5.46 (d, *J* = 9.2 Hz, 1H), 5.32 (dd, *J* = 16.6, 5.3 Hz, 2H), 5.16 (d, *J* = 5.4 Hz, 1H), 4.61 (t, *J* = 5.5 Hz, 1H), 3.73 – 3.67 (m, 2H), 3.44 – 3.37 (m, 3H), 3.23 – 3.19 (m, 1H), 2.69 (dt, *J* = 27.3, 7.8 Hz, 4H), 2.11 (s, 3H), 1.77 (p, *J* = 7.9 Hz, 2H).

^13^C NMR (126 MHz, DMSO-*d*_6_) δ 184.6, 184.1, 146.4, 146.1, 143.3, 133.8, 131.7, 131.6, 125.9, 125.8, 121.0, 87.4, 79.9, 77.0, 72.2, 69.6, 60.8, 27.9, 26.3, 25.2, 12.5.

HRMS (ESI-TOF, *m/z*) for C_22_H_26_O_7_ 444.1693 [M + H]^+^, found 444.2313.

#### 2.3.9 2-methyl-3-(3-(1-(2-(((2*R*,3*R*,4*S*,5*S*,6*R*)-3,4,5-trihydroxy-6-(hydroxymethyl)tetrahydro-2*H*-pyran-2-yl)oxy)ethyl)-1*H*-1,2,3-triazol-4-yl)propyl)naphthalene-1,4-dione (Trojan Horse 4)

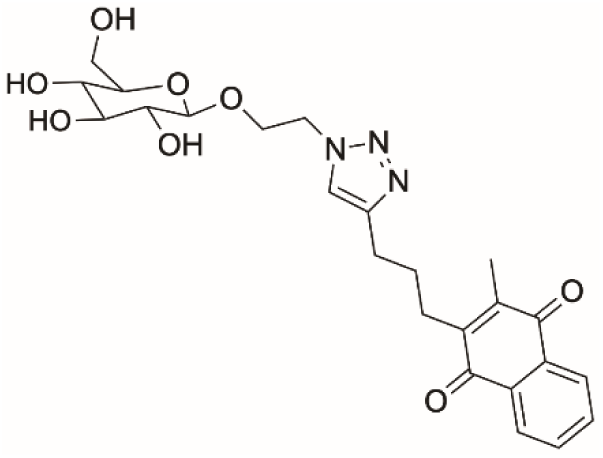

Prepared according to the General CuAAC procedure from alkyne **5** (86 mg, 0.361 mmol) and 2-azidoethyl β-D-glucopyranoside (75 mg, 0.301 mmol) (3 h, 100% CH_2_Cl_2_ to 3% MeOH in CH_2_Cl_2_) to afford the title compound as a yellow solid (89 mg, 61%).

1H NMR (500 MHz, DMSO-*d*_6_) δ 8.02 – 7.98 (m, 2H), 7.95 (s, 1H), 7.85 – 7.81 (m, 2H), 4.54 – 4.46 (m, 4H), 4.21 (d, *J* = 7.8 Hz, 1H), 4.07 – 4.03 (m, 1H), 3.87 – 3.83 (m, 1H), 3.67 (d, *J* = 11.8 Hz, 1H), 3.42 (dd, *J* = 11.7, 6.0 Hz, 2H), 3.17 – 3.10 (m, 3H), 3.02 (t, *J* = 9.1 Hz, 1H), 2.96 (t, *J* = 8.4 Hz, 1H), 2.70 (t, *J* = 7.5 Hz, 2H), 2.64 (t, *J* = 7.8 Hz, 2H), 2.09 (s, 3H), 1.77 (p, *J* = 7.7 Hz, 2H).

^13^C NMR (126 MHz, DMSO-*d*_6_) δ 184.6, 184.1, 146.2, 146.1, 143.2, 133.8, 131.7, 131.6, 125.9, 125.8, 122.7, 102.9, 77.0, 76.6, 73.3, 70.0, 67.4, 61.1, 49.4, 27.8, 26.2, 25.2, 12.5.

HRMS (ESI-TOF, *m/z*) for C_24_H_30_N_3_O_3_ 488.1955 [M + H]^+^, found 488.1591.

### 2.4 Cytotoxicity and Selectivity

Cells were plated in a 96-well black fluorescent plate (Fisher Scientific, Ireland) at a density of 20 x 10^4^ in complete media and placed in varying concentrations of glucose media together with the compound under investigation, which was dissolved in 10 µL of sterile water. A 20 % DMSO solution served as a cell death control to ensure 100 % cell death. The cells were incubated for either 2, 24 or 48 h.

Following treatment alamar blue assay was conducted as per supplier’s instructions. Plates were refrigerated overnight at 4 °C, and then their fluorescent emission at 604 nm (578 nm excitation) was recorded on the GloMax microplate reader (Promega, Madison WI, USA). The values obtained were normalised against the blank well readings and used to plot the dose response sigmoidal curves to determine the IC_50_ values.

Statistical analyses were performed using GraphPad Prism version 9.0 (GraphPad Software, San Diego, CA, USA) and Microsoft Excel (Microsoft Corporation, Redmond, WA, USA). Fluorescent emission values obtained from the alamar blue assay were normalized to blank well readings to correct for background fluorescence. Dose-response curves and IC50 values were calculated using nonlinear regression analysis with a four-parameter logistic model in GraphPad Prism. Data from the cytotoxicity assays were analysed using one-way ANOVA followed by Dunnett’s multiple comparisons test to compare treated groups with control groups. A p-value of < 0.05 was considered statistically significant.

The selectivity index (SI) measures a compound’s cytotoxic activity in cancer cells compared to non-malignant cells and is calculated as a ratio of IC_50_ (non-malignant): IC_50_ (cancer). An SI greater than 1.0 indicates that the compound is more effective against cancer cells compared to non-malignant cells, whereas an SI less than 1.0 suggests a non-selective response with similar efficacy in both cell types.

### 2.5 Metabolic Phenotype and Bioenergetics

The Seahorse MitoStress Test evaluates critical aspects of mitochondrial function, providing insights into cellular energy demands and mitochondrial health. Basal respiration indicates the cell’s energetic demand under baseline conditions and is directly linked to ATP production. ATP production measures the portion of oxygen consumption specifically used for synthesising ATP. Proton leak represents the amount of oxygen consumption not coupled to ATP production, serving as an indicator of mitochondrial membrane integrity. Maximal respiration assesses the cell’s maximal respiratory capacity, while spare respiratory capacity, defined as the difference between maximal and basal respiration, indicates the cell’s ability to respond to increased energy demand. Non-mitochondrial respiration accounts for the oxygen consumption that is not linked to mitochondrial activity. Together, these parameters offer a comprehensive picture of mitochondrial function and cellular metabolism.

The Seahorse ATP Rate Test measures key parameters related to ATP production where the rate of ATP synthesis is quantified from both glycolysis and oxidative phosphorylation respectively, the two primary pathways for cellular energy production. It distinguishes the contributions of each pathways to ATP production and assesses the cell’s reliance on mitochondrial respiration versus glycolysis. This comprehensive analysis reveals shifts in metabolic phenotype under various conditions, helping to understand cellular energy balance, identify metabolic reprogramming, and evaluate the effects of treatments on cellular bioenergetics.

#### 2.5.1 MitoStress Test

Cells were seeded at 6 x 10^4^ cells/well in a 24-well XF microplate (Agilent Technologies, Santa Clara, CA, USA). Specific inhibitors and uncouplers were prepared in XF assay media, which was supplemented with 0 mM D-(+)-glucose for sequential addition at the appropriate final concentrations of oligomycin A (1.8 µM), FCCP (4 µM) and antimycin A (2 µM) (all Sigma-Aldrich). Cells were placed in a non-CO_2_ incubator at 37 °C for 1 h prior to the assay. Basal respiration (OCR) and extracellular acidification rate (ECAR) were measured by the Seahorse Biosciences XFe24 Extracellular Flux Analyser (Agilent Technologies, Santa Clara, CA, USA). All recorded measurements were normalised to cell number, determined using the crystal violet assay.

#### 2.5.2 ATP Rate Test

Inhibitors and uncouplers were prepared in XF assay media, supplemented with 0 mM D-(+)-glucose for sequential addition at the appropriate final concentrations of oligomycin A (1.8 µM), and antimycin A (2 µM) (all Sigma-Aldrich). Cells were placed in a non-CO_2_ incubator at 37 °C for 1 h prior to the assay. Basal respiration (OCR) and extracellular acidification rate (ECAR) were measured by the Seahorse Biosciences XFe24 Extracellular Flux Analyser (Agilent Technologies, Santa Clara, CA, USA).

MitoStress and ATP Rate Tests: For the MitoStress and ATP Rate tests, oxygen consumption rates (OCR) and extracellular acidification rates (ECAR) were measured and analysed using the Agilent online analysis tool. Baseline readings were established, and the effects of specific inhibitors and uncouplers on OCR and ECAR were assessed using repeated measures ANOVA and plotted as bar charts, and scatter plots. Error bars indicating standard deviation were included to illustrate variability within the data.

### 2.6 ROS and MMP Flow cytometry

Cells were seeded at 6 x 10^4^ cells per well in a 96-well plate and incubated for 24 h. Fluorescent dyes were then added to the plate and incubated for 30 mins at room temperature, protected from light, according to the supplier’s instructions. Cells were all resuspended in 60 µL of FACS. Compensation was performed with positive and negative compensation beads. Gating and analysis were performed using CellStream Analysis software.

Flow cytometry data for ROS and mitochondrial membrane potential (MMP) were analysed with CellStream Analysis software (Luminex Corporation, Austin, TX, USA).

Compensation for the flow cytometry was performed with positive and negative compensation beads, and gating was set based on control samples. Mean fluorescence intensity (MFI) was compared between treated and control groups using unpaired t-tests. The data presented was represented by fold change scatter plots.

Fold change was calculated by dividing the value of the parameter in treated cells by the value in untreated cells. A fold change greater than 1 indicates an increase in the parameter, while a fold change less than 1 indicates a decrease.

### 2.8 Metabolomics

Metabolomic analysis was performed using Liquid Chromatography (LC) tandem Mass Spectrum (MS) with the QTRAP 6500 LC-MS/MS system, equipped with a multi-component Ion Drive. This advanced method leverages SCIEX patented Ion Drive technology to enhance ion production, transmission, and detection, thereby ensuring high sensitivity and performance. The system offers improved polarity switching and multiple reaction monitoring (MRM) speeds, facilitating expedited chromatography and enhanced throughput. The integrated QTRAP enables simultaneous quantitative MRMs and qualitative QTRAP scans in a single injection, maximizing efficiency.

#### 2.8.1 Sample Preparation

PNT1a, LNCaP, PC3, and Du145 cells were seeded at 5 x 10^6^ cells/mL in T75 flasks and incubated for 24 hr. Following incubation, cells were washed with DPBS, and then incubated in 8 mL of RPMI assay medium containing the desired glucose concentration (0 – 11 mM glucose) (Sigma Aldrich), supplemented with 10% dialyzed FBS (Sigma Aldrich). Vitamin solutions, prepared at IC50 concentrations of Menadione, Trojan Horse 4, and Trojan Horse 6 compounds, were added to respective flasks. After another 24 hr of incubation, cells were gently trypsinised, centrifuged, washed with PBS, resuspended, and flash-frozen in liquid nitrogen. The samples were stored at -80 °C until they were sent to the University College Dublin (UCD) Metabolomics core facility for processing and analysis.

#### 2.8.2 Metabolomics Analysis

Sample preparation followed the MxP® Quant 500 assay manual (Biocrates Life Sciences, Innsbruck, Austria). The analytical procedures involved derivatization, drying, addition of high-performance liquid chromatography (HPLC)-grade water, and subsequent LC-MS/MS and flow injection analysis tandem mass spectrometry (FIA-MS/MS) analyses.

#### 2.8.3 Data Processing and Metabolite Quantification

Data processing and metabolite quantification were performed using MetIDQ software provided by Biocrates Life Sciences. Amino acids, amino acid-related metabolites, and biogenic amines were quantified based on isotopically labelled internal standards and calibration curves. Semi-quantification of other metabolites utilized internal standards. Data quality assessment included analysis of accuracy and reproducibility of quality control samples.

#### 2.8.4 Data and Statistical Analysis of Metabolite Concentrations

Two hundred and eleven metabolites were detected above the limit of detection (LOD) from the five hundred examined. Statistical analysis was performed using a two-tailed t-test between groups of interest (p ≤ 0.05). Significant metabolites were further analysed using repeated measures (Bonferroni) 1-way ANOVA with multiple comparisons applied.

## 3.0 Results

### 3.1 Synthesis of the menadione-glucose and fatty acid conjugated compounds by click chemistry

The menadione-sugar and fatty acid conjugates were synthesised using a Cu(I)-catalyzed azide-alkyne cycloaddition (CuAAC) between azido derivatives of glucose and a fatty acid to two alkyne menadione compounds (Figure 1). A CuAAC strategy, commonly known as a “click” reaction, was chosen as the synthesis provides facile access to triazoles from the appropriate azides and alkynes and this linking group is considered a privileged bioisostere in drug discovery programs.^35^

**Figure 1.**
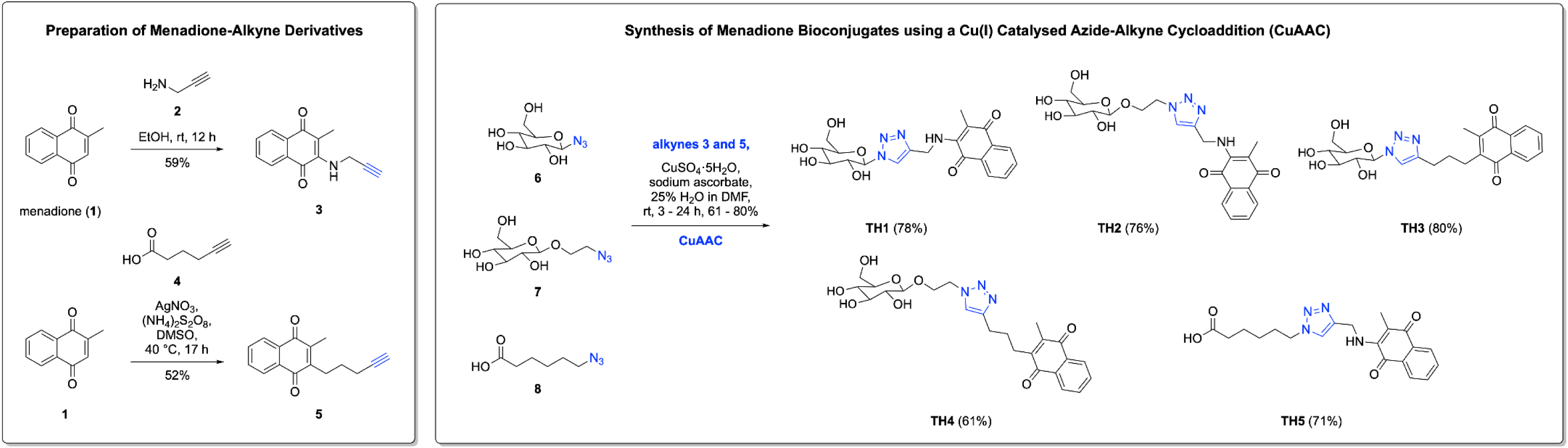
Synthesis of menadione-based alkynes **3** and **5** via conjugate addition followed by a spontaneous oxidation and alkylation via a silver-catalysed decarboxylation. Alkynes **3** and **5** were then “clicked” to β-D-glucopyranosyl azide **6**, 2-azidoethyl β-D-glucopyranoside **7** and 6-azido-hexanoic acid **8** to give triazoles TH1-5 in excellent yields of 61 to 80%.

Menadione **1** was functionalised with reactive alkyne handles following literature procedures to give the amino **3** and alkyl **5** derivatives in synthetically useful yields of in 59% and 52%, respectively.^33^ The alkynes were then “clicked” to the azido sugars **6** and **7**, and fatty acid **8** to furnish the triazoles TH1 – 5 in 61 – 80% yields (Figure 1).

### 3.2 Activity in Human Drug-Resistant Prostate Cancer Cells

Five menadione-glucose and menadione-fatty acid compounds were synthesised by copper-catalysed click chemistry via an azide or amine linker group as seen in Figure 2 (a). These modifications to the metabolic substrates used were with the goal to increase the selectivity towards human cancer cells due to the unique metabolism displayed in PCa.

**Figure 2:**
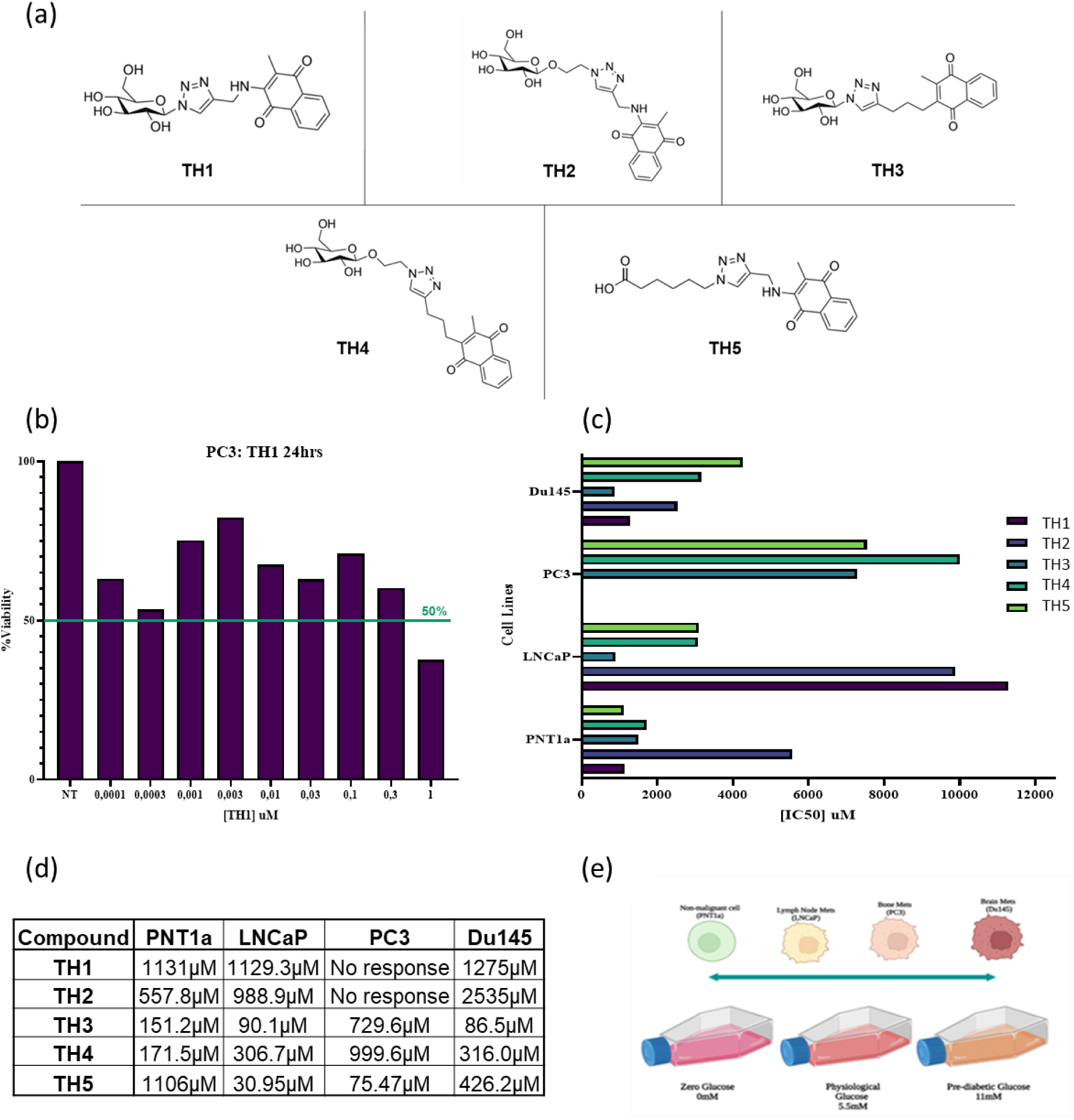
**(a)** Molecular structures of the Trojan Horse (TH) compounds investigated in the study. TH1-4 are composed of Menadione with a sugar backbone, while TH5 consists of menadione with a fatty acid (6-azidohexanoic acid) backbone. **(b)** Initial assessment of TH1 revealed poor cytotoxicity in the presence of media glucose (11mM). **(c)**.IC_50_ values of compounds TH2-6 in all 4 cell lines examined in the preliminary screening for the compound efficacy. **(d)** Screening of the first six compounds across all four cell lines identified TH3 (glucose) and TH5 (fatty acid) as the optimal compounds for further investigation in the study. **(e)** A glucose milieu was employed to assess the competitive nature of glucose-conjugated compounds and glucose present in the cellular medium

Preliminary cytotoxicity analysis was carried out on TH1 compound in the PC3 cells under media glucose conditions (11 mM) where there was very little cell killing, and the IC_50_ values were high, as shown in Figure 2 (b) This suggested that perhaps due to the glucose molecule conjugated to Menadione, the uptake of the compound was being inhibited and therefore it prompted us to consider the impact of the glucose milieu on the uptake of the TH compounds and the role of media glucose in modulating cytotoxicity. Therefore varying glucose conditions were examined to represent glucose starvation (0 mM), normal physiological glucose (5.5 mM), and high glucose (11 mM) conditions.

### 3.3 Screening for Cytotoxicity and Selectivity in Human Prostate Cancer Cells

An examination of the cytotoxicity and selectivity of the novel menadione-glucose/fatty acid compounds were performed on a panel of prostate non-malignant and cancer cell lines across 3 glucose conditions. The cell lines included PNT1a (prostate non-malignant), LNCaP (androgen dependent), PC3 and Du145 (androgen independent) under zero glucose, 5.5 mM glucose and 11 mM glucose media conditions. While all the cell lines showed some toxicity towards the 3 compounds of interest (TH1, TH3 and TH5), PNT1a showed no cytotoxicity towards TH1 up to 1000 µM concentrations. The compounds TH3 (glucose) and TH5 (lipid) presented a similar cytotoxicity towards the cell lines, but with an IC_50_ of ∼300µM obtained across the three glucose conditions when treated with TH5.

The selectivity was determined for each drug under each glucose conditions in each cell line, with an optimal selectivity index (SI) deemed to be greater than 1.0. TH5 had a high SI of 3.1 in the DU145 cells at zero glucose and an overall SI of ≥1 across the cell lines and glucose conditions (Table 1 (b)). TH3 had a SI of 2.0 in the DU145 cells in the 5.5 mM glucose medium, with overall higher SI values than TH1 (Table 1 (b)).

**Table 1.** **(a)** The numeric values represent the calculated IC50 cytotoxicity of each compound in the four cell lines under three glucose conditions following a 24-hour treatment period. All values are expressed in micromolar concentrations (µM). This data provides insight into the cytotoxic efficacy of the compounds across various cellular environments, facilitating comparative analysis and interpretation of treatment outcomes. **(b)** The selectivity index (SI) represents the ratio of cytotoxicity observed in cancer cell lines to that observed in non-malignant cell lines under specific treatment and glucose conditions. A higher SI indicates greater cytotoxicity towards cancer cells relative to non-malignant cells. The selectivity was determined for each drug under each glucose condition in each cell line. In this study an optimal SI was defined as greater than 1. This index provides insights into the relative cytotoxicity of the drugs, specifically comparing their effects on cancer cells to those on non-malignant cells under identical treatment and glucose conditions.

| (a) IC50 Concentrations |  |  |  |  |  |  |  |  |  |
| --- | --- | --- | --- | --- | --- | --- | --- | --- | --- |
| Cell line | zero Glucose |  |  | 5.5mM Glucose |  |  | 11mM Glucose |  |  |
|  | TH1 IC50 (μM) | TH3 IC50 (μM) | TH5 IC50 (μM) | TH1 IC50 (μM) | TH3 IC50 (μM) | TH5 IC50 (μM) | TH1 IC50 (μM) | TH3 IC50 (μM) | TH5 IC50 (μM) |
| PNT1a | 14.4 | 27.7 | 92.8 | 148.0 | 323.8 | 341.0 | 284.5 | 338.2 | 315.2 |
| LNCaP | 339.8 | 82.4 | 297.7 | 368.8 | 79.6 | 295.7 | 377.5 | 86.6 | 344.9 |
| PC3 | N/A | 317.5 | 113.8 | N/A | 1537.0 | 326.8 | N/A | 2085.0 | 309.9 |
| Du145 | 37.7 | 103.6 | 29.9 | 241.1 | 162.8 | 305.9 | 214.5 | 241.6 | 317.2 |

**Table 1 (a)** The numeric values represent the calculated IC50 cytotoxicity of each compound in the four cell lines under three glucose conditions following a 24-hour treatment period.
| (b) Selectivity Index |  |  |  |  |  |  |  |  |  |
| --- | --- | --- | --- | --- | --- | --- | --- | --- | --- |
| Cell line | zero Glucose |  |  | 5.5mM Glucose |  |  | 11mM Glucose |  |  |
|  | TH1 | TH3 | TH5 | TH1 | TH3 | TH5 | TH1 | TH3 | TH5 |
| LNCaP | 0.04 | 0.3 | 0.3 | 0.4 | 1.8 | 1.2 | 0.8 | 1.8 | 0.9 |
| PC3 | N/A | 0.09 | 0.8 | N/A | 0.2 | 1 | N/A | 0.2 | 1 |
| Du145 | 0.3 | 0.3 | 3.1 | 0.6 | 2 | 1.1 | 1.3 | 1.4 | 0.99 |

### 3.4 Effect on Prostate Cancer Cell Metabolic Phenotypes and Bioenergetics

An examination of the effects of the novel TH compounds on the mitochondrial bioenergetics as well as the metabolic phenotype was conducted. All 3 TH analogues were examined in parallel with inhibitors and modulators of cellular metabolic pathways. The oxygen consumption rate (OCR) and extracellular acidification rate (ECAR) were measured to determine changes in mitochondrial respiration and glycolysis, respectively, following treatment with TH1, TH3, and TH5 under zero, 5.5mM and 11mM glucose conditions.

Treatment with TH5 led to a reduction in glycolytic activity, as indicated by decreased rate of glycolysis and ECAR, specifically in the non-malignant PNT1a cells (Figure 3). This reduction was most evident in the zero glucose condition, suggesting a potential impairment in glycolytic flux when glucose availability is restricted. In contrast, the cancer cell lines LNCaP, PC3, and Du145 did not show significant changes in glycolysis under the same conditions, indicating a possible adaptive response that maintains glycolytic activity even in the presence of TH5.

**Figure 3.**
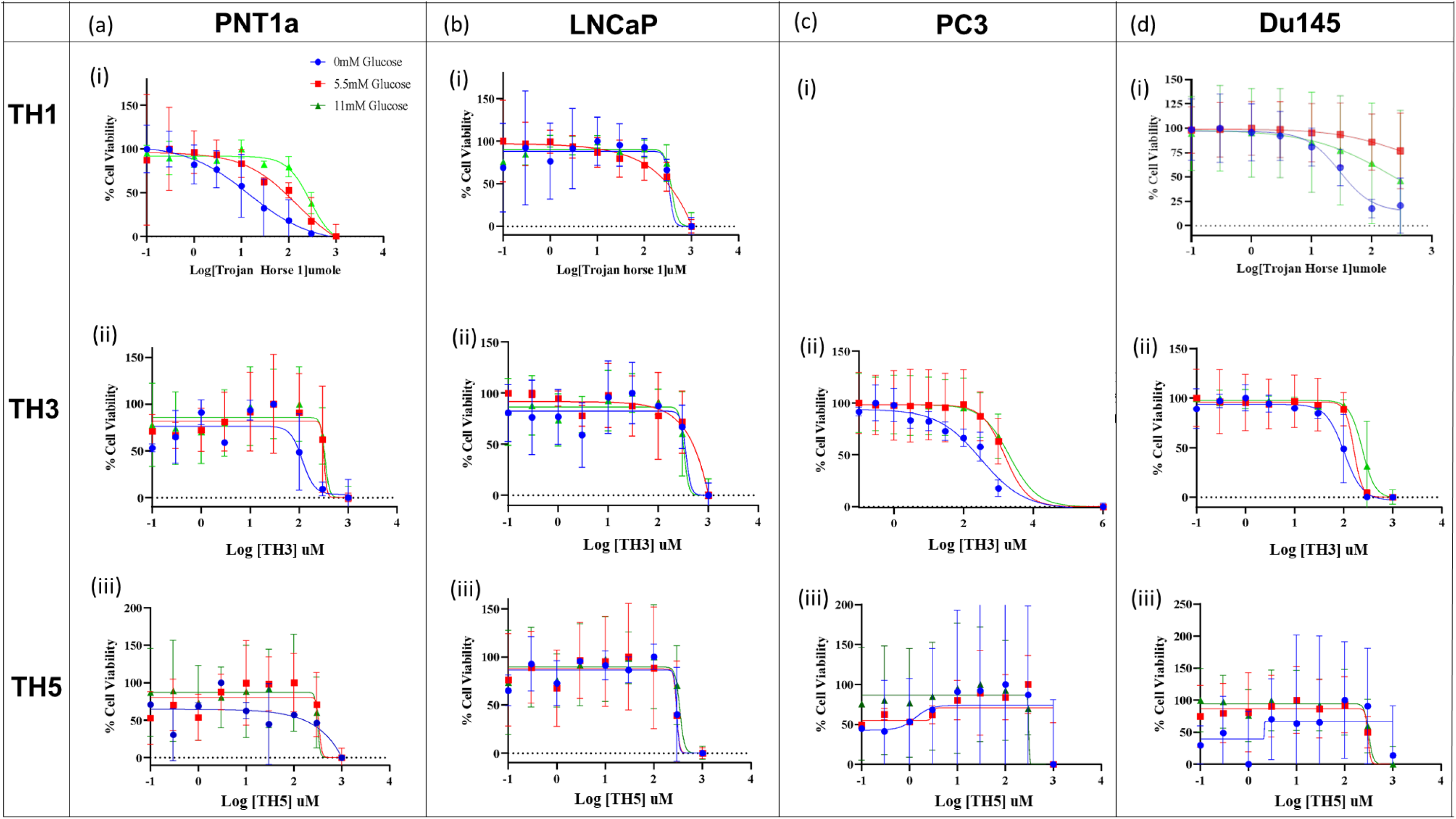
Cytotoxic effects of TH compounds on prostate cell lines under varying glucose concentrations after 24hrs. **(a)** PNT1a: (i) TH1, (ii) TH3, (iii) TH5. **(b**) LNCaP: (i) TH1, (ii) TH3, (iii) TH5. **(c)** PC3: (i) TH1 (no response), (ii) TH3, (iii) TH5. **(d)** Du145: (i) TH1, (ii) TH3, (iii) TH5. Cells were plated at a density of 2 x 10^4 cells/well in 96-well plates and treated with menadione derivatives at various concentrations for 24 h. Cell viability was assessed using the alamar blue assay. Dose-response curves were generated to determine IC50 values. Data are presented as mean ± SD from three independent experiments performed in triplicate. Blue denotes zero glucose conditions, red signifies 5.5 mM glucose, and green indicates 11 mM glucose. **(a)** PNT1a cells. This figure demonstrates the cytotoxic effects of the novel TH compounds across different prostate cell lines, highlighting the variability in response based on androgen dependency, metastatic origin, and glucose concentration.

With regards to OxPhos, the treatment with TH5 resulted in a reduction of OCR across all prostate cancer cell lines, with the most significant decrease observed in LNCaP cells in 5.5 mM glucose conditions (Figure 4 b(ii)). This suggests that TH5 impacts mitochondrial respiration more profoundly in androgen-dependent versus the androgen-independent cell lines. Additionally, the observed reduction in maximal respiration and proton leak in TH5 treated cells further supports the hypothesis that TH5 may compromise mitochondrial function by inhibiting oxidative phosphorylation through the high production of ROS, leading to a decrease in overall mitochondrial activity (Figure 5).

**Figure 4:**
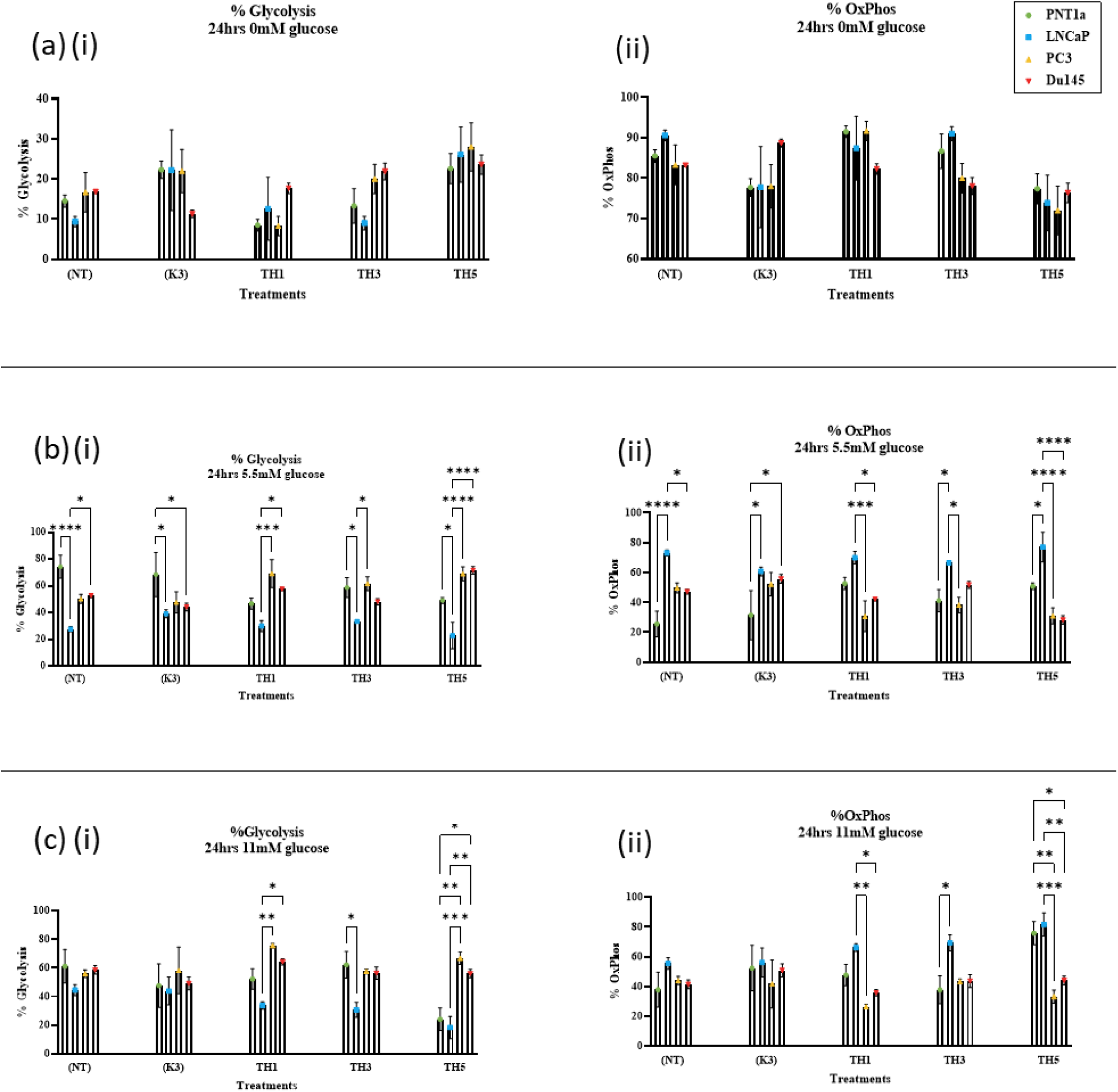
ATP production in prostate cell lines under varying glucose conditions and treatments **(a)** Zero Glucose ATP production. (i) Percentage of Glycolysis: PNT1a (green), LNCaP (blue), PC3 (yellow), and Du145 (red) cells under zero glucose conditions, with untreated control, native menadione, TH1, TH3, and TH5 treatments. (ii) Percentage of Oxidative Phosphorylation (OxPhos) **(b)** 5.5 mM Glucose ATP production. (i) Percentage of Glycolysis. 5.5 mM glucose conditions. Elevated glycolysis rates are observed in all cell lines except LNCaP. (ii) Percentage of OxPhos. **(c)** 11 mM Glucose ATP production. (i) Percentage of Glycolysis under 11 mM glucose conditions. (ii) Percentage of OxPhos. Data represent mean ± SEM from three independent experiments performed in triplicate. Statistical analysis was conducted using one-way ANOVA (*p<0.05).

**Figure 5.**
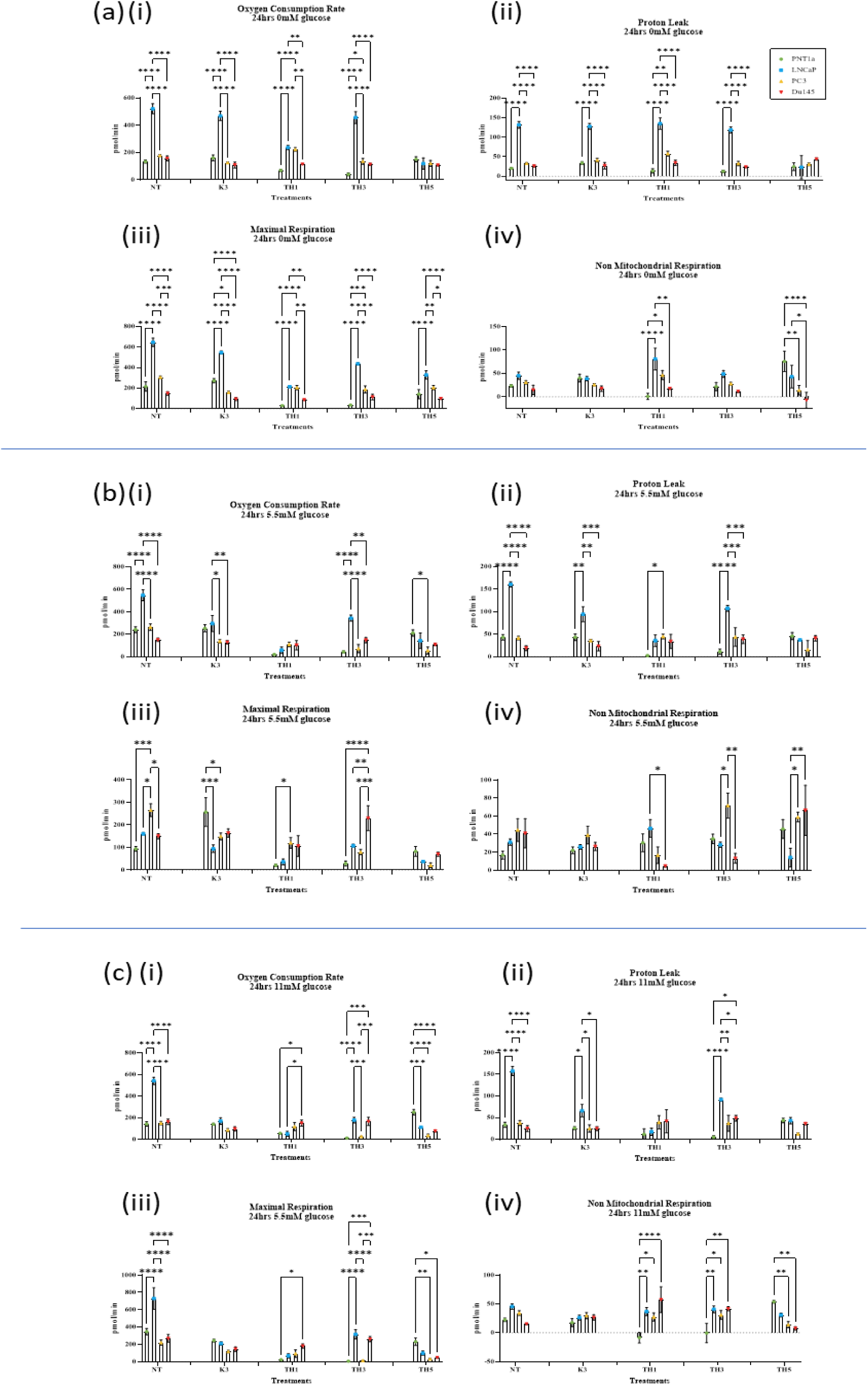
Mitochondrial bioenergetics endpoints in prostate cell lines under varying glucose conditions and treatments. Mitochondrial bioenergetics endpoints in PNT1a (green), LNCaP (blue), PC3 (yellow), and Du145 (red) cells. Treatments include Untreated control (NT), menadione control (K3), TH1, TH3, and TH5. (a) Zero Glucose Conditions. (b) 5.5 mM Glucose Conditions. (c) 11 mM Glucose Conditions. (i) Oxygen Consumption Rate (OCR). (ii) Proton Leak. (iii) Maximal Respiration. (iv) Non-mitochondrial Respiration. Data represent mean ± SD from three independent experiments performed in triplicate. Statistical analysis was conducted using two-way ANOVA (*p<0.05).

Notably, TH1 and TH3 treatments did not produce significant alterations in either OCR (% OxPhos) or ECAR (%glycolysis), highlighting a distinct difference in the efficacy of these compounds relative to TH5. These results suggest that TH5 may be more effective at disrupting both glycolytic and mitochondrial pathways by the proposed production of ROS. in certain cell lines, particularly PNT1a and LNCaP.

### 3.5 Effect on Prostate Cancer Cell ROS, Mitochondrial Membrane Potential and Apoptosis

The effects of the TH compounds on cell death, ROS production, and mitochondrial membrane potential (MMP) were assessed via flow cytometry.

The results indicated a variable response across different prostate cell lines. Specifically, in non-malignant PNT1a cells, treatment with TH3 and TH5 led to a statistically significant increase in ROS levels (Figure 6 a). In contrast, this ROS increase was not observed in the prostate cancer cell lines LNCaP, PC3, and Du145. The heightened ROS levels in PNT1a suggest that these non-malignant cells have a lower capacity to upregulate antioxidant defences such as superoxide dismutase (SOD), catalase, and glutathione peroxidase, which are typically elevated in cancer cells, as part of their metabolic reprogramming.^36^

**Figure 6:**
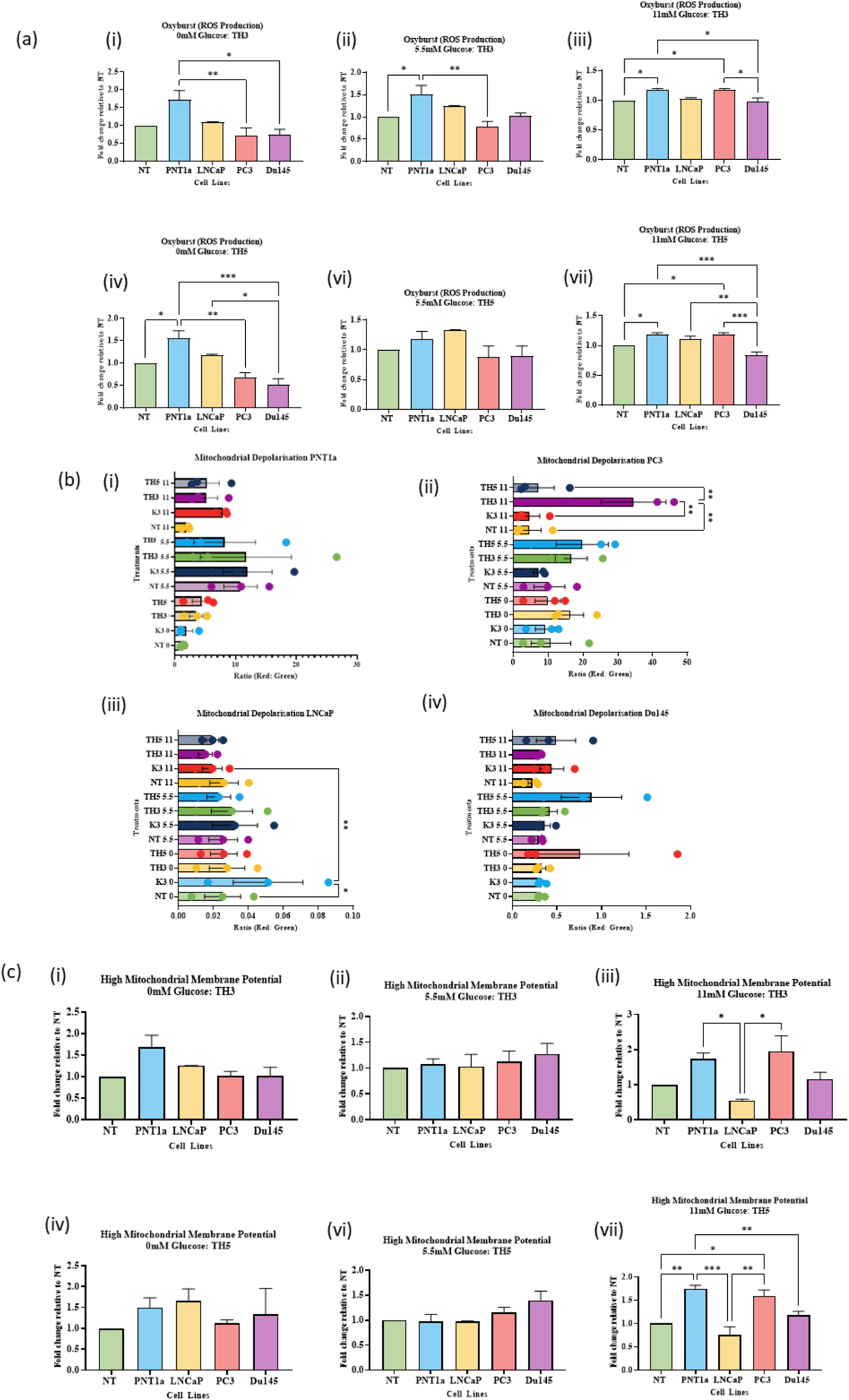
Fold Change Analysis of Reactive Oxygen Species (ROS) and Mitochondrial Membrane Potential (MMP) in Prostate Cell Lines. This figure illustrates the relative differences in ROS and MMP levels between treated and untreated prostate cell lines under varying glucose conditions. **(a)** Reactive Oxygen Species (ROS) Fold Change. **(b)** Mitochondrial Membrane Potential (MMP) Fold Change. (i) Zero Glucose TH3 treated. (ii) 5.5 mM Glucose TH3 treated. (iii) 11 mM Glucose TH3 treated. (iv) Zero Glucose TH5. (v) 5.5 mM Glucose TH5. (vi) 11 mM Glucose TH5. Data represent mean ± SEM from three independent experiments performed in triplicate. Statistical analysis was conducted using two-way ANOVA (*p<0.05).

Contrary to our hypothesis that TH treatment would evoke widespread ROS-induced mitochondrial damage and subsequent cell death, ROS levels in cancer cells remained unchanged. This stability in ROS levels might indicate the cancer cells’ ability to counteract ROS through enhanced antioxidant mechanisms. This was supported by the observation that MMP levels, indicative of mitochondrial health, remained stable across all cell lines and treatment conditions (Figure 6 c), suggesting that mitochondrial integrity was maintained despite TH compound treatment.

Additionally, mitochondrial depolarization (Figure 6 b), a precursor to apoptosis, was evaluated under varying glucose conditions across all cell lines treated with TH compounds. However, only small changes were detected (Figure 6 b i-ii). Despite the lack of change in ROS levels and MMP in cancer cell lines, apoptosis was still observed, particularly in the TH5 treated groups, also seen in the cytotoxic evaluations in Figure 3.

### 3.6 Metabolomic determination of ROS scavenging Amino Acids

Untargeted metabolomic screening of cell pellets from the 3 glucose conditions and treatments with TH3, TH5 revealed significant alterations in 33 metabolites, including 21 amino acids, 5 biogenic acids, 3 carboxylic acids, and 4 ceramides. Pathway mapping using Metaboanalyst 5.0 identified metabolites linked to Warburg Metabolism, such as lactic acid, which was elevated in PNT1a, PC3, and Du145 cells, particularly in the 11 mM glucose TH5 treatments. This lactic acid accumulation, indicative of Warburg glycolysis, is associated with late-stage metastatic prostate cancer and poor prognosis, due to its role in promoting metastasis, angiogenesis, and immunosuppression, shown in Figure 9 of the supplemental section.

Several amino acids like cysteine, carnosine, and proline, known for their roles in oxidative stress management, were increased in various treated cells. Elevated cystine levels, particularly in Menadione-treated PNT1a cells, suggest an increased demand for ROS scavengers induced by treatment. Similarly, increased carnosine levels in LNCaP and PC3 cells treated with TH5 may be a response to elevated endogenous ROS levels. Proline elevation in TH-treated LNCaP and PC3 cells may indicate ROS scavenging activity to counteract Menadione-induced ROS generation. Methionine elevation in TH5-treated PNT1a cells suggests a role in regulating ROS levels or redox homeostasis. These alterations underscore the adaptive antioxidant responses of cancer cells to cope with ROS influx, potentially contributing to their survival against TH treatments.

The PCA biplot (Figure 7a) illustrates the correlation of amino acid metabolomic data from four prostate cell lines (PNT1a, LNCaP, PC3, and Du145) under the various TH treatment groups and glucose conditions. PNT1a cells exhibit a clear separation based on glucose availability, those in the presence of glucose (5.5mM and 11mM) cluster tightly in the left quadrant, while samples in zero glucose conditions form a distinct cluster in the centre-right of the plot. LNCaP cells cluster together in the lower-centre right of the plot, showing little variation between glucose starved and the presence of glucose conditions. PC3 cells show a separation between samples in the presence of glucose (centre cluster) and those in zero glucose conditions (above centre cluster), indicating a distinct clustering pattern based on glucose availability. Du145 cells, like LNCaP, show no clear separation based on glucose conditions and instead cluster centrally.

**Figure 7:**
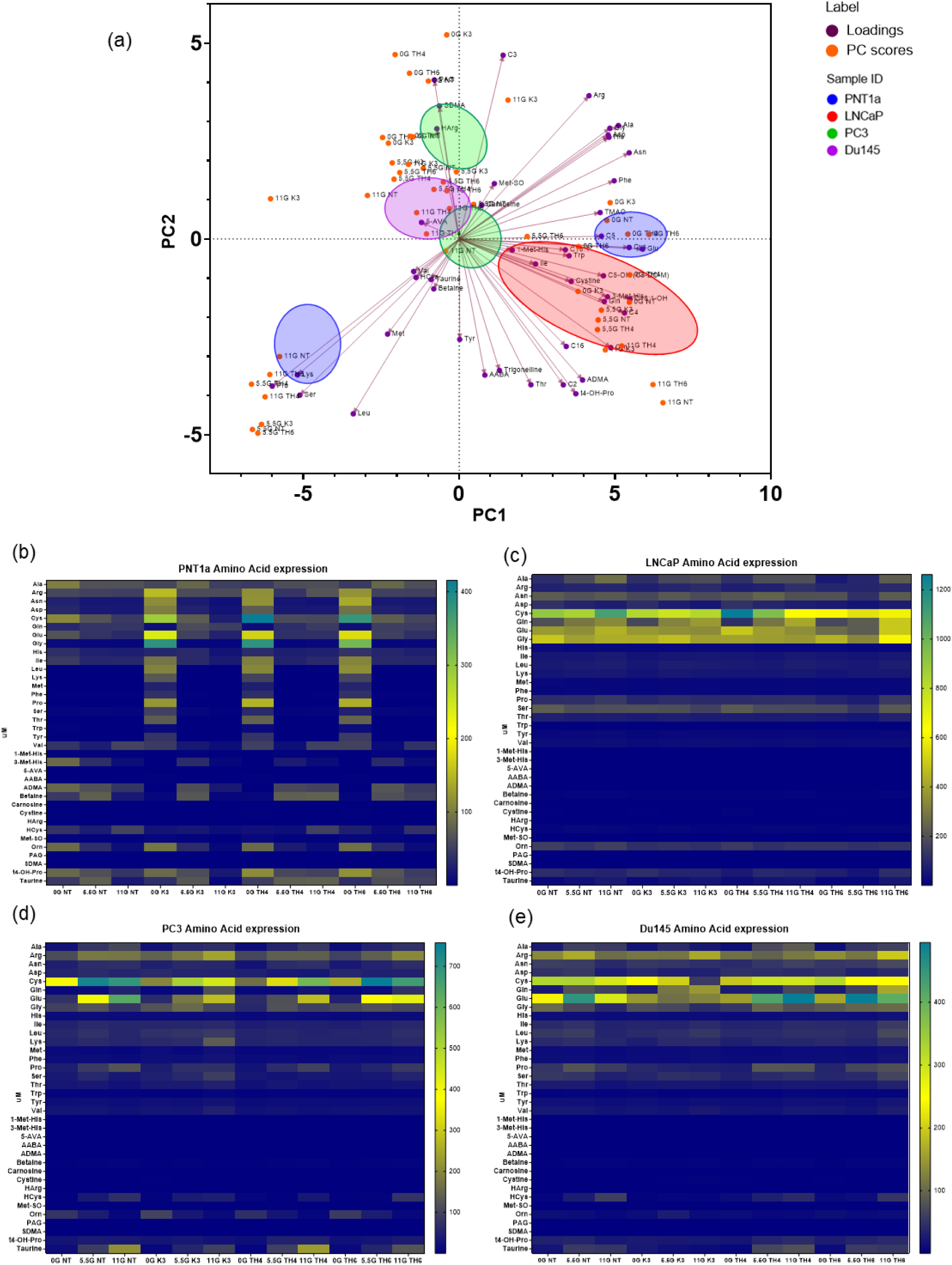
Amino Acid Metabolomic Analysis in Prostate Cell Lines **(a)** The PCA biplot illustrates the correlation of amino acid metabolomic data, where the loadings are scaled by a multiplier to allow the Principal Component (PC) scores and loadings to be displayed on the same graph. PCA analysis identifies patterns in the data by reducing its dimensionality, revealing how different amino acids contribute to variations across treatment groups and glucose conditions. The biplot shows the relationship between cell lines, treatments and glucose conditions with the amino acids, highlighting clusters and trends in metabolite expression. Correlation clustering is denoted by blue PNT1a, red is LNCaP, green is PC3 and purple is Du145**. (b)** Heatmap representation of amino acid levels in PNT1a cells across different treatment groups and glucose conditions. Dark green shading indicates higher, yellow is intermediate and navy is low metabolite expression, providing a visual comparison of the relative abundance of each amino acid under various experimental conditions. (c)-(e) represent the same data but for the other cell lines evaluated. **(c)** LNCap **(d)** PC3 **(e)** Du145. Data represent mean values from three independent experiments performed in triplicate. The PCA biplot and heatmaps provide a comprehensive overview of the impact of treatments and glucose conditions on amino acid metabolite levels in different prostate cell lines. Statistical analysis was conducted using appropriate multivariate methods to identify significant differences and correlations (*p<0.05). This figure demonstrates the distinct metabolic profiles of prostate cell lines in response to different treatments and glucose conditions, highlighting the variability in amino acid metabolite expression.

Across all cell lines, there is no clear separation based on TH treatment groups within each cell type, suggesting that the primary drivers of variation in the PCA biplot are cell type and glucose condition rather than the specific TH treatment applied.

### 4.0 Discussion

Prostate cancer is one of the most common cancers among men worldwide, making advancements in understanding its pathogenesis and potential treatments crucial for reducing the global healthcare burden and improving patient outcomes. PCa cells have an altered metabolic phenotype, with early disease relying on OxPhos, and more intermediate/late disease switching to fatty acid oxidation and Warburg metabolism.^22,37^ Gaining a better understanding of the PCa metabolic processes could provide novel targets for treatment. In this study we aimed to examine various prototype compounds that facilitate the uptake of compounds that limit the metabolic capacity of PCa cells. These TH compounds comprised of a basic fuel molecule conjugated to a non-toxic vitamin moiety, to target the cancers unrelenting need for fuel.

Our hypothesis for the TH compounds anticipated that the malignant cells would have a higher uptake of the glucose conjugated TH compounds compared to Menadione. This expectation is grounded in the well-established principle of the Warburg effect, which states that malignant cells consume glucose at a rate approximately 100 times faster than non-malignant cell.^38,39,40^ Menadione has been extensively studied both in vitro and in vivo for its anticancer effects, both alone and in combination with conventional cancer therapies, showing promising result and high tolerance (in *vivo*).^41–44^

Two methods of conjugation were employed to attach Menadione with the metabolic moiety-either a sugar or a lipid-with the goal of enhancing the selective cytotoxicity of these compounds in cancer cells while minimising their effects on normal cells. Among these, the best cytotoxicity and selectivity was seen in the TH3 and TH5 compounds, though the SI values remained relatively low. Of the glucose-Menadione conjugates, TH3 exhibited a higher SI than TH1 across various cell lines and glucose conditions. The difference between TH1 and TH3 lies in their conjugation groups: TH1 features an amine linker, whereas TH3 has an aryl linker chain. This variation in chemistry allowed TH3 to show greater selectivity towards metastatic cells compared to non-malignant cells, highlighting the importance of not only the active elements (Menadione and glucose) but also the efficacy of the linker groups.

In click chemistry, limits are placed on the linker group composition, suggesting the need for other methods of compound conjugation in therapeutic synthesis. The highest SI determined was 3.1 with the TH5 treatment in the metastatic Du145 cells. For new cancer therapeutics to be considered suitable for clinical efficacy, a high SI is essential. While an Si of 1 is deemed selective, an SI of 10.0 and higher is often required for safety in *vivo*.^45^ The SI values obtained for the TH compounds in this study indicate that further development is needed to reach the optimal SI values for safe in vivo application. Given menadione’s success in existing research, modifications to the linker group and the metabolic substrate may enhance the overall selectivity of the compounds.^44,46,47^

Treatment resistance in cancer remains a significant issue, with many studies investigating strategies to overcome cellular mechanisms of resistance. The key factor in this resistance is the plasticity of cancer which allow them to shift to a differentiated state, with limited tumorigenic potential thereby supporting their continued growth and proliferation.^48,49^ This study aimed to target this plasticity by developing a treatment that targets multiple aspects of cancer cell nutrient uptake, specifically focussing on the Warburg effect. Ultimately, the goal was to trigger redox related cell death through the cellular uptake of menadione. Although some efforts have aimed to disrupt the metastatic process by glycoconjugation of natural substrates and known chemotherapies, these approaches have not been successful to date.^50,51^

We postulated that treatment with the TH compounds could induce a metabolic switching event within the PCa cells, reversing Warburg metabolism by reducing glycolysis, and thereby regressing the disease. However, this shift was not observed in the TH treated cells. Instead, our findings highlighted that androgen independent metastatic PCa cells remained persistently engaged in the Warburg effect, even after nutrient deprivation and the novel treatments. This highlights the importance of the Warburg effect in late stage PCa.

Research by Pertega-Gomes et al has showed the metabolic heterogeneity of PCa and its clinical relevance, illustrating that the advanced stages of PCa, both in *vitro*, and in *vivo,* present increased glycolytic activity, which correlates with poorer patient prognosis.^52^Their work showed increase glucose consumption in PC3 cells along with increased OCR and ECAR overall highlighting their metabolic plasticity.^52^ This raises the question of whether it is possible to reverse the Warburg phenotype in androgen independent cells and restore the metabolic programming observed in androgen dependent LNCaP cells, potentially preventing treatment resistance in PCa. With this we aimed to investigate how varying glucose conditions affect the metabolism of PCa cell lines and whether our TH compounds could alter their metastatic potential.

Although no significant changes were observed in the metabolic phenotypes of the cells when treated with the TH compounds, alterations were observed in the mitochondrial bioenergetics. The cells treated with the novel compounds across the glucose conditions presented at times with increased proton leak, maximal respiration, basal OCR, and non-mitochondrial respiration all linked to alterations in the mitochondria and possible mitochondrial dysfunction. Mitochondrial health was further examined through the MMP and mitochondrial depolarisation. The link between mitochondrial bioenergetics and MMP are noted frequently in the literature, with increased mitochondrial depolarisation and a sustained shift in MMP indicating mitochondrial dysfunction overall impacting OxPhos in the mitochondria, then impacting the cells OCR. A decrease in mitochondrial depolarisation was not found in any of the cell lines in the study, however alterations in MMP was found in the cells treated with menadione and TH5. Although, the androgen independent metastatic disease (Du145 cells) did not present with any alterations to their MMP or mitochondrial depolarisation. However, the alterations that were observed in the mitochondrial bioenergetics were anticipated to likely be due to heightened ROS levels in the cells due to the treatment with the novel compounds.^53^

We hypothesized that mitochondrial dysfunction seen in the bioenergetic profiles of the cell lines was due to an increase in endogenous ROS due to Menadione induced oxidative stress. Menadione is thought to reduce oncogenic superoxide levels, leading to apoptosis via the generation of onco-suppressive hydroperoxides and cytotoxic hydroxyl radicals.^11,14^ ROS scavengers and antioxidants quench ROS, decreasing the oxidative stress capabilities of menadione overall reducing its anticancer effects.^53–56^ However, if high enough concentrations of Menadione are achieved, the capability of antioxidant enzymes to eliminate ROS is exceeded and results in redox related death.^53–56^ The results of the ROS measurements were surprising, as only small increases in ROS were observed in the cancer cells treated with Menadione and the TH compounds, contrary to our initial expectations. However, the metabolomic analysis showed increased levels of ROS scavenging amino acids, such as cysteine, carosine and proline in cells treated with the novel compounds. Cysteine, carnosine, and proline are all amino acids linked to oxidative stress management and are known ROS scavengers increased in times of oxidative stress.^57–59^ The three amino acids were found in the PNT1a, LNCaP and PC3 cells treated with the novel compounds, which may account for the lack of ROS determined through the Oxyburst assay by flow cytometry. Betaine and methionine are also implicated in redox homeostasis, the influx in these amino acids may remove excess ROS produced by the TH compounds. The metabolomic evaluation of the cell lines tied all the pieces together giving light as to why high levels of ROS was not observed in the cells treated with the TH compounds. The amino acids are creating a dynamic balance between the ROS produced by the compounds and the cell’s ability to produce ROS scavenging molecules to maintain ROS homeostasis. Surprisingly, we did still observe metabolic alterations in the cells through the cellular bioenergetics even with ROS scavenging observed which may be an effort by the cells to alter their metabolism to increase amino acid production as a protective effect against ROS. From this, the metabolic reprogramming of the disease back to its earlier disease phase may be possible with therapeutic intervention, however there are many biological modulators that must be considered.

The treatment with TH compounds led to an increase in ROS scavengers, which in turn affected amino acid metabolite expression. Studies have shown that androgen can activate amino acid metabolism in PCa. It would be worthwhile to investigate whether this treatment also alters AR expression in the cell lines, as this could impact metabolic alterations within the cell.^60,61^ Putluri et al found an influx? (or enhancement?) in pathways associated with amino acid metabolism in androgen treated PCa cells, implicating that androgen signalling plays a role in metabolic regulation.^60^ Again, with an interest in reversing the androgen independent metastatic phenotype by perturbing the Warburg effect, this may also impact the androgen status of the cancer. Many chemotherapeutics of PCa target androgens, so reversing this phenotype reversal could have huge implications on the clinical outcomes for patients with currently untreatable PCa.^62^

Existing studies have shown that the PC3 cell line resembles CRPC and is a suitable model for intermediate androgen-independent disease.^63–65^ When PC3 were incubated with 11mM glucose and treated with TH3, an increase in ROS was seen. This finding was encouraging as the IC_50_ values required to induce cell death in PC3 cells were significantly higher compared to other cell lines. This indicated that the levels of TH compounds required to cause an increase in ROS production is far higher than previously thought. Additionally, PC3 cells treated with the TH compounds, exhibited elevated levels of ROS scavenging amino acids, which might explain the lower-than-anticipated ROS levels.

In conclusion, our study demonstrates the intricate relationship between metabolic reprogramming and therapeutic interventions in PCa. Although we faced challenges in perturbing the Warburg effect in androgen-independent PCa cells, our exploration of novel metabolic-targeting compounds shows promise for advancing therapeutic strategies. Future work should focus on refining these compounds to enhance selectivity and reduce cytotoxicity in non-malignant cells. Understanding PCa metabolism and its impact on disease progression and therapy response is crucial. Continued investigation into the mechanisms of metabolic reprogramming and androgen signalling could offer new insights into disease management and treatment efficacy. Ultimately, reversing the PCa disease phenotype with future compound iterations could significantly improve patient outcomes and reduce the burden of untreatable disease. In essence, our study contributes to the ongoing efforts to unravel the intricacies of PCa metabolism and developing innovative therapeutic interventions. By leveraging our understanding of metabolic dysregulation in cancer cells, we aspire to pave the way for personalised and targeted treatments that enhance patient outcomes.

## Author contributions

Initial Concept: DAB, RDB, SS, and JOL. Project Conceptualization, JJOL, DAB, PT, LBE, RDB, EFH and SS. Data curation: LBE, FOC,PT, RDB and EFH. Formal cellular analysis, LBE, FOC. Funding acquisition, JJOL, DAB, RDB. Investigation, LBE, PT, FOC, EFH, and RDB. Methodology and synthesis, LBE, PT, FOC, EFH and RDB. Project administration: PT, SO, CM, DAB, RDB, JJOL. Visualization, LBE. Writing—original draft based on PhD thesis, LBE. Review and editing—all authors have reviewed, edited and agreed to publish this version of the manuscript.

## Acknowledgements

A component of the funding for this project was provided by Envision Sciences Pty Ltd. DAB is a founding shareholder and DAB and RDB have benefited from Envision Sciences Pty Ltd.’s research funding. JJOL is a shareholder and has benefited from Envision Sciences Pty Ltd.’s research funding. Graphical Abstract created with BioRender.com.

## Conflict of Interest

The authors declare no conflicts of interest.

## Declaration of Transparency and Scientific Rigour

This Declaration acknowledges that this paper adheres to the principles for transparent reporting and scientific rigour of preclinical research as stated in the *BJP* guidelines and as recommended by funding agencies, publishers and other organisations engaged with supporting research.

## Abbreviations

AAT: Androgen Ablation Therapy
AIPC: Androgen Independent Prostate Cancer
ECAR: Extracellular Acidification Rate
MMP: Mitochondrial Membrane Potential
OCR: Oxygen Consumption Rate
OxPhos: Oxidative Phosphorylation
PCa: Prostate Cancer
ROS: Reactive Oxygen Species
SI: Selectivity Index
SOD: Superoxide Dismutase
TH: Trojan Horse

## Supplemental Figures

**Figure 9:**
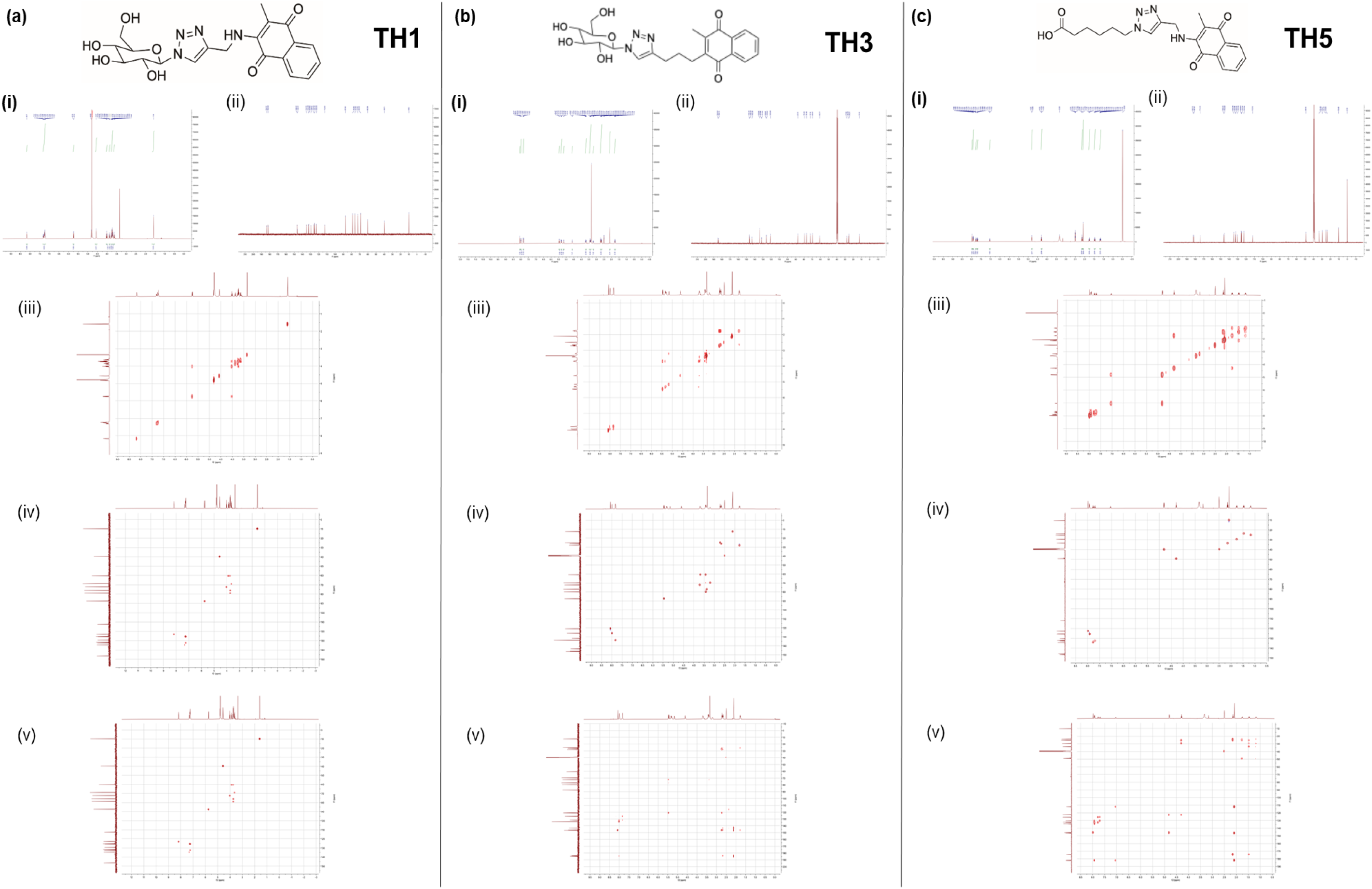
Structural spectral analysis of the lead compounds of the study. (a) Trojan Horse 1 (TH1), (b) Trojan Horse 3 (TH3) and (c) Trojan Horse 5 (TH5). (i) ^1^H NMR (ii) ^13^C NMR (iii) COSY NMR (iv) HSQC NMR (v) HMBC NMR. Full spectral analysis of all compounds available.

**Figure 10:**
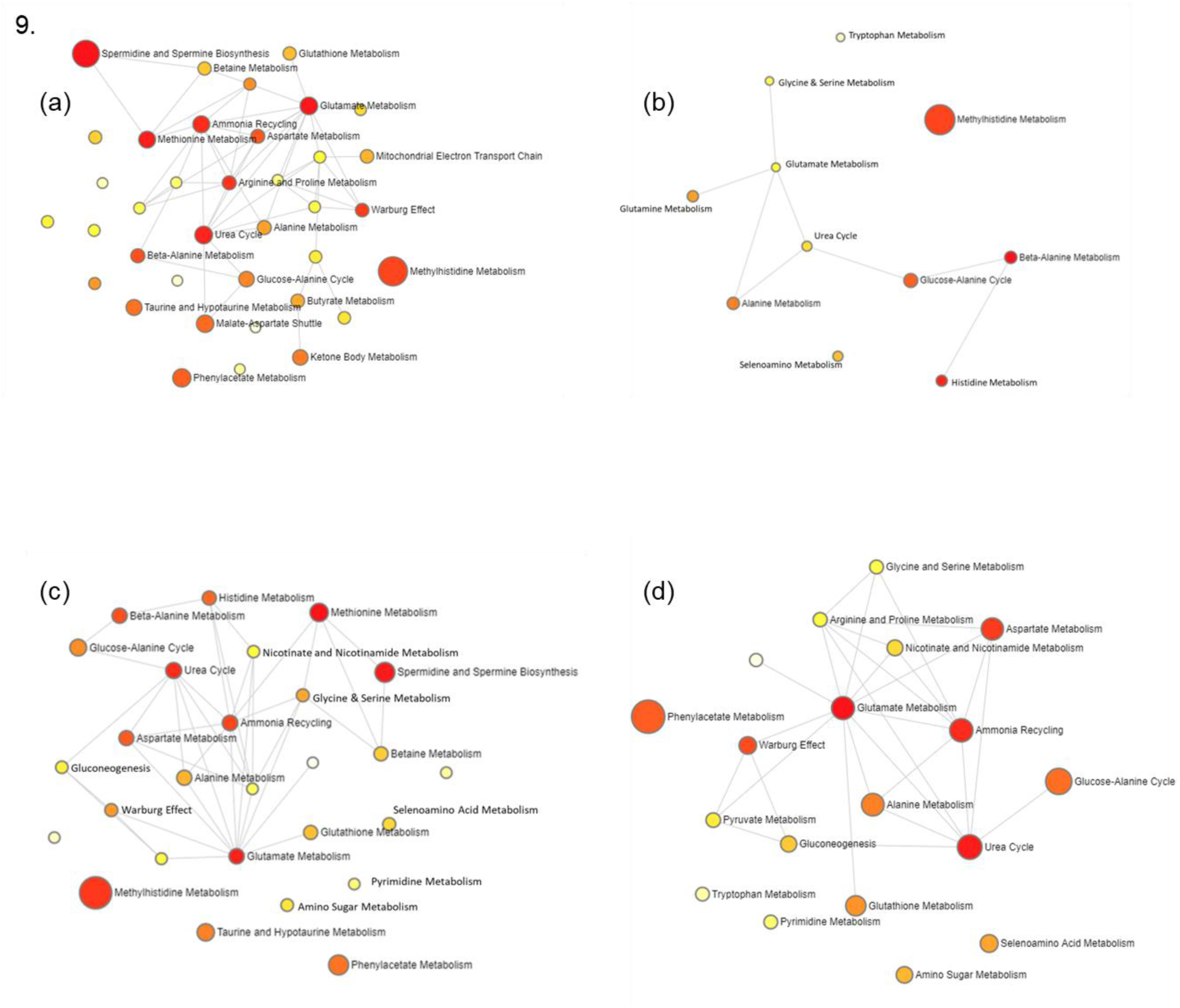
Pathway analysis of the significant metabolites present in the cell lines (a) PNT1a, (b) LNCaP, (c) PC3 and (d) Du145; indicating the possible pathways of relevance in the cell line. Small yellow dots indicate pathways of less significance, with the larger red dots indicating pathways of higher significance. Connected dots indicate pathway connections. Pathways were determined by the number of metabolites present within each pathway based on the human metabolome database (HMDB) and matched to metabolic pathways by MetaboAnalyst 5.0.

## Notes

### Competing Interest Statement

The authors have declared no competing interest.

